# Interleukin-19 alleviates experimental autoimmune encephalomyelitis by attenuating antigen-presenting cell activation

**DOI:** 10.1101/2020.07.15.204826

**Authors:** Hiroshi Horiuchi, Bijay Parajuli, Hiroyasu Komiya, Yuki Ogawa, Shijie Jin, Keita Takahashi, Yasu-Taka Azuma, Fumiaki Tanaka, Akio Suzumura, Hideyuki Takeuchi

## Abstract

Interleukin-19 (IL-19) acts as an anti-inflammatory cytokine in various inflammatory diseases. Multiple sclerosis (MS) is a major neuroinflammatory disease in the central nervous system, but it remains uncertain how IL-19 contributes to MS pathogenesis. Here, we demonstrate that IL-19 deficiency aggravates experimental autoimmune encephalomyelitis (EAE), a mouse model of MS, by promoting IL-17–producing helper T cell (Th17 cell) infiltration into the central nervous system. In addition, IL-19–deficient splenic macrophages expressed elevated levels of major histocompatibility complex class II, co-stimulatory molecules, and Th17 cell differentiation–associated cytokines such as IL-1β, IL-6, IL-23, TGF-β1, and TNF-α. These observations indicated that IL-19 plays a critical role in suppression of MS pathogenesis by inhibiting macrophage antigen presentation, Th17 cell expansion, and subsequent inflammatory responses. Furthermore, treatment with IL-19 significantly abrogated EAE. Our data suggest that IL-19 could provide significant therapeutic benefits in patients with MS.

## Introduction

Multiple sclerosis (MS) and experimental autoimmune encephalomyelitis (EAE), a mouse model of MS, are major autoimmune demyelinating diseases of the central nervous system (CNS)^1,2^. Various types of immune cells and soluble mediators contribute to the complex mechanisms underlying the onset and progression of both MS and EAE, and recent studies have shown that type 1 helper T (Th1) cells and interleukin-17–producing helper T (Th17) cells play pivotal roles in their pathogenesis^3, 4, 5^. In these diseases, autoreactive Th17 cells primed in the lymph nodes infiltrate the CNS and activate microglia/macrophages that induce inflammatory demyelination and subsequent neuronal damage, resulting in a wide range of clinical features, including sensory and motor paralysis, blindness, pain, incontinence, and dementia^1, 2^.

Interleukin-19 (IL-19) is an IL-10 family cytokine that is homologous and highly similar to IL-20 and IL-24^6, 7^. IL-19 binds to the heterodimeric receptor consisting of IL-20Rα and IL-20Rβ, and its downstream signaling is mediated by STAT3 phosphorylation^8, 9^. IL-19 is mainly produced by activated macrophages and microglia^10, 11, 12, 13^. Recent studies showed that IL-19 exerts anti-inflammatory effects on macrophages by inhibiting inflammatory cytokine production, downregulating antigen-presenting capacity, and enhancing M2 phenotype polarization, which promotes type 2 helper T (Th2) cell differentiation and suppresses Th1 and Th17 cell differentiation^11, 12, 14, 15, 16, 17^. In fact, IL-19 plays a critical role in development of various autoimmune diseases, including asthma^18, 19^, psoriasis^20, 21, 22^, inflammatory bowel disease^11, 23^, rheumatoid arthritis^24^, and Type I diabetes^25^. However, it remains to be elucidated how IL-19 contributes to MS pathogenesis.

Here, we examined the pathological role of IL-19 in EAE using IL-19–deficient (IL-19^−/−^) mice. IL-19 deficiency markedly exacerbated EAE, and treatment with IL-19 effectively suppressed EAE accompanied by inhibiting macrophage antigen presentation and subsequent expansion of Th17 cells. Our findings suggest that IL-19 may provide significant therapeutic benefits for treating MS.

## Results

### IL-19 deficiency exacerbates EAE

We generated myelin oligodendrocyte glycoprotein (MOG)-induced EAE in C57BL/6J wild-type (WT) and IL-19–deficient (IL-19^−/−^) mice. IL-19^−/−^ mice exhibited earlier disease onset and more severe symptoms than WT mice (Fig. 1A). Histological analysis of the lumbar spinal cords revealed more inflammatory cell infiltration in IL-19^−/−^ mice than in WT mice (Fig. 1B). Flow cytometric analysis also disclosed that IL-19^−/−^ EAE mice had more CNS-infiltrating cells than WT EAE mice (Fig. 1C). We then chronologically evaluated IL-19 expression levels in the spleen and lumbar spinal cord of WT mice at pre-immunization, disease onset, and disease peak. Splenic IL-19 mRNA expression was upregulated at disease onset, but was strongly suppressed at the disease peak (Fig. 1D). By contrast, upregulation of IL-19 mRNA in the lumbar spinal cord was observed at disease peak (Fig. 1E). These results suggest that endogenous IL-19 serves as a negative regulator of EAE pathogenesis at both the induction and effector phases.

**Figure 1.**
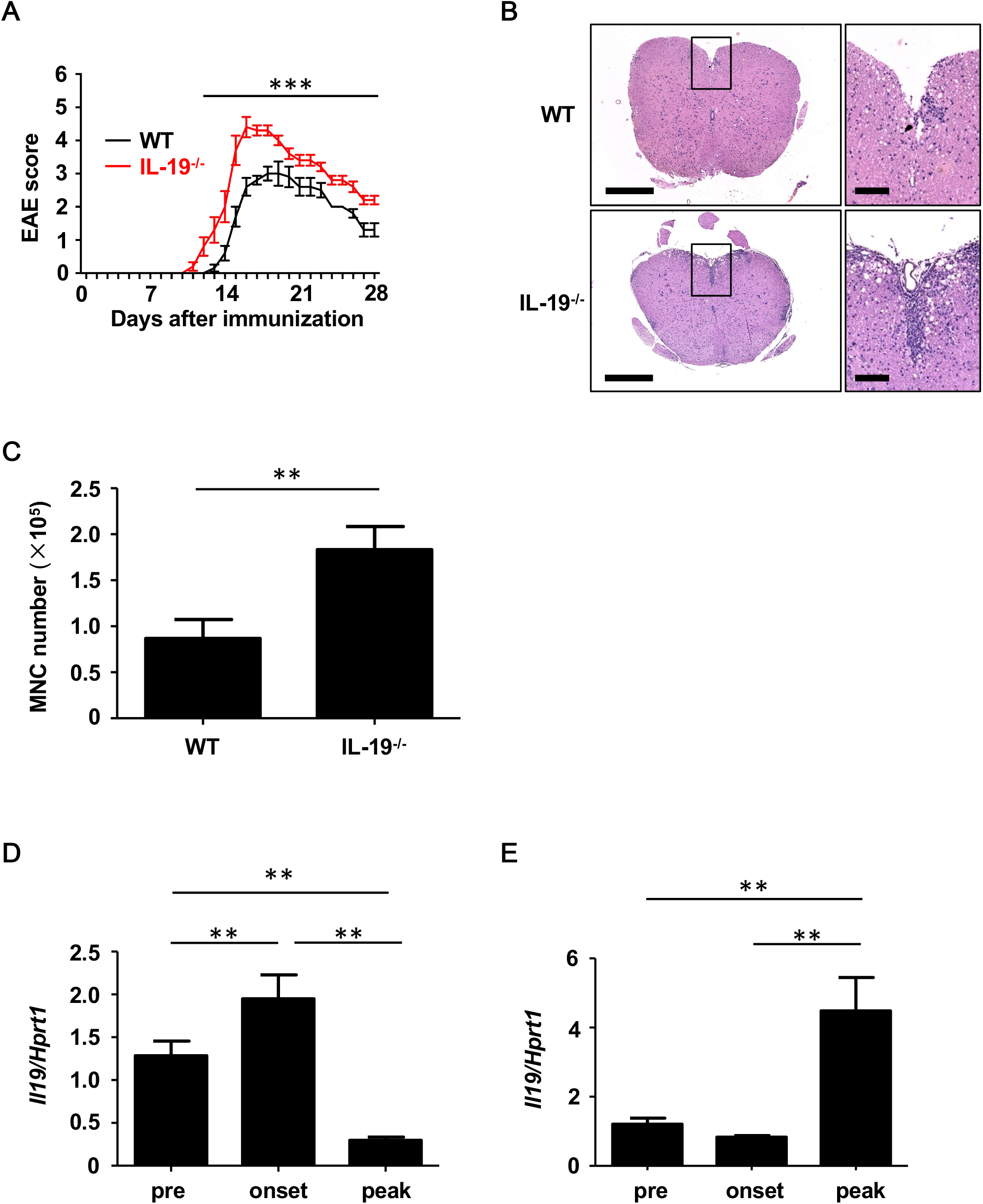
IL-19 deficiency aggravates EAE. (A) EAE clinical scores for WT (black) and IL-19^−/−^ (red) mice. Data represent means ± SD (n = 10). ***, *p* < 0.0001. (B) Micrographs of hematoxylin/eosin staining of L5 lumbar spinal cords at the peak EAE of WT and IL-19^−/−^ mice. The right panels show enlargements of the boxed areas in the left panels. Scale bars: 500 μm (left), 100 μm (right). (C) Flow cytometric analysis of cell infiltration in the CNS at the peak EAE (n = 5). Data represent means ± SD. **, *p* < 0.01. (D and E) chronological qPCR data for IL-19 mRNA expression level in the spleen (D) and the lumbar spinal cord (E) of WT EAE mice (n = 5). Pre, pre-immunization; onset, EAE onset; peak, EAE peak. Data represent means ± SD. **, *p* < 0.01.

### IL-19 deficiency increases Th17 cell infiltration into the CNS

Because EAE is a Th1 and Th17 cell–mediated autoimmune disease, we next assessed whether IL-19 deficiency would increase CNS infiltration of Th1 and Th17 cells. Flow cytometric analysis revealed that at disease peak, IL-19^−/−^ mice exhibited more infiltration of Th17 cells in the spinal cord than WT mice (Fig. 2A, B). By contrast, no difference was observed in Th1 cell infiltration between IL-19^−/−^ and WT mice (Fig. 2C). These results indicate that IL-19 deficiency mediates elevated CNS infiltration by Th17 cells, but not Th1 cells.

**Figure 2.**
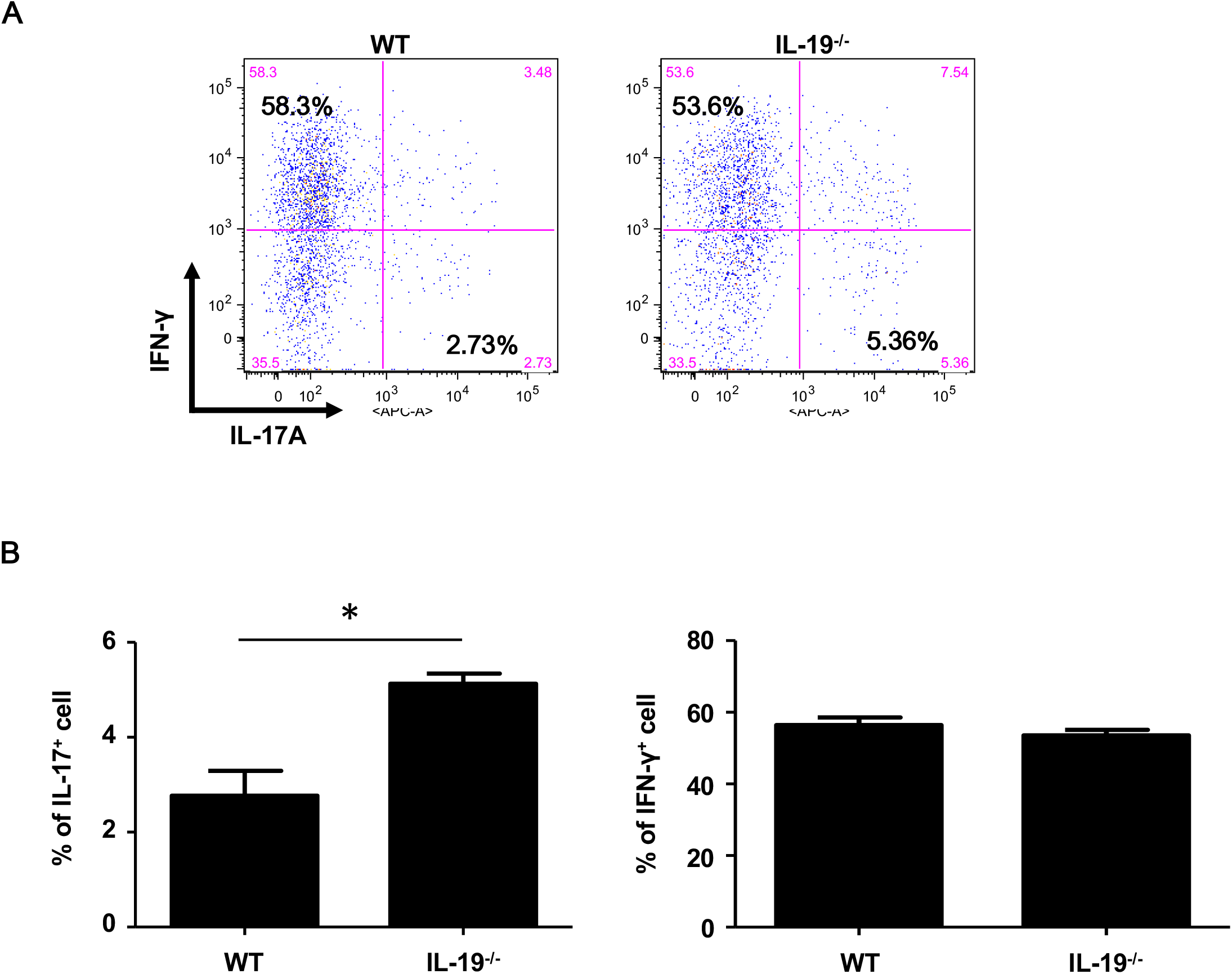
IL-19 deficiency increases CNS infiltration of Th17 cells, but not Th1 cells. (A) Representative flow-cytometric data for IL-17– and IFN-γ–producing CD4^+^ T cells in the CNS at the peak EAE. (B) Percentage of IL-17–producing CD4^+^ T cells. (C) Percentage of IFN-γ–producing CD4^+^ T cells. Data represent means ± SD (n = 3). *, *p* < 0.05.

### IL-19 deficiency expands Th17 cell population

To determine whether IL-19 contributes to the expansion of Th17 cells during the induction phase of EAE, we assessed the antigen-specific expansion of Th17 cells *ex vivo*. Splenic CD4^+^ T cells isolated from WT and IL-19^−/−^ mice at MOG-EAE onset were stimulated with MOG peptide for 3 days *in vitro*. IL-19 deficiency significantly upregulated *Il17a* mRNA and downregulated *Foxp3* mRNA, but it did not affect the level of *Ifng* mRNA (Fig. 3A). Flow cytometric analysis also revealed that IL-19 deficiency expanded the Th17 cell population (Fig. 3B, C). These results indicate that IL-19 deficiency mediates expansion of Th17 cells in the peripheral lymphoid tissues.

**Figure 3.**
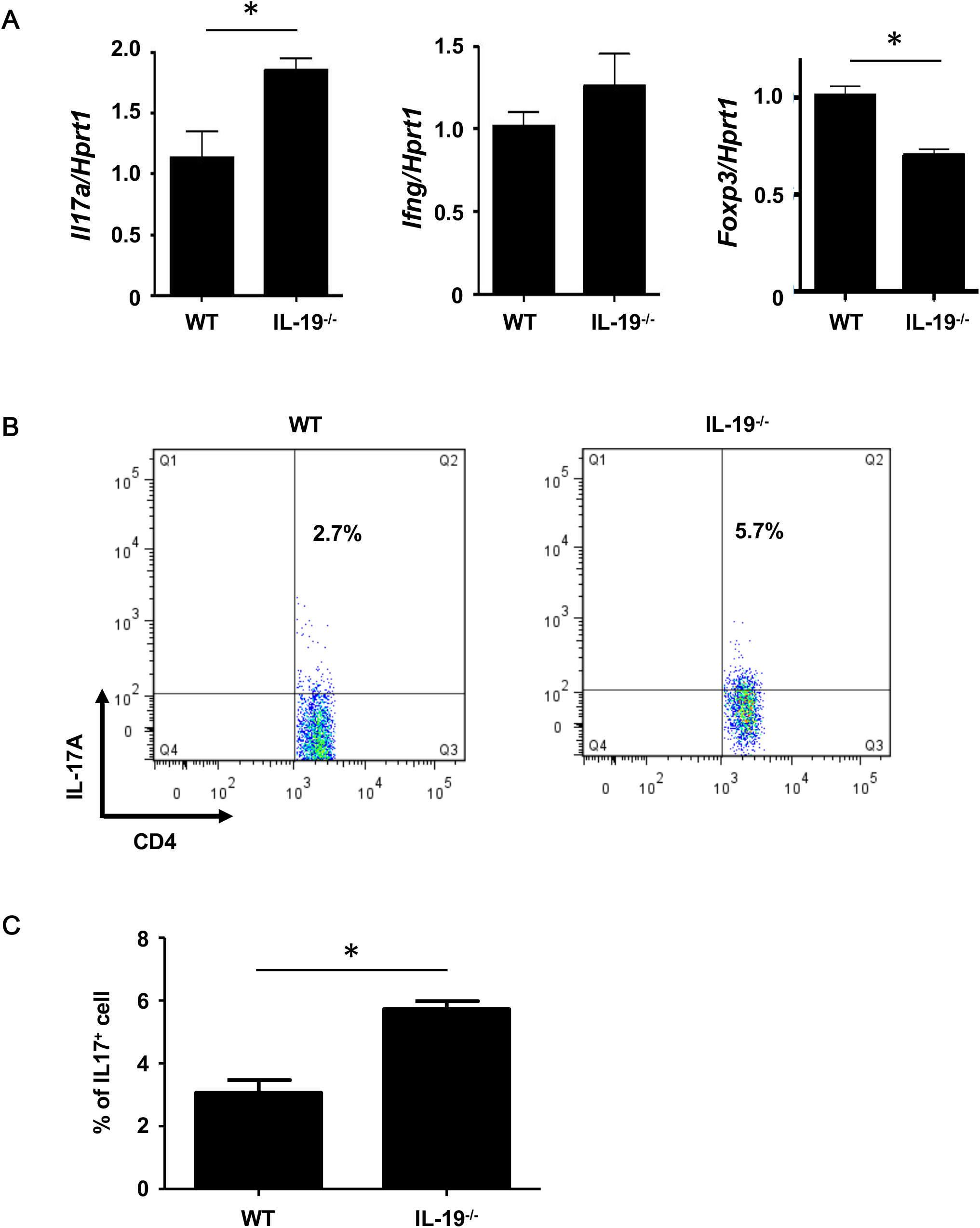
IL-19 deficiency expands the Th17 cell population. (A) qPCR data for levels of mRNAs encoding IL-17A, IFN-γ, and FoxP3 in splenic CD4^+^ T cells. (B) Representative flow cytometric data for IL-17–producing CD4^+^ T cells in the spleens of EAE mice. (C) Percentage of IL-17–producing CD4+ T cells in the spleens of EAE mice. Data represent means ± SD (n = 3). *, *p* < 0.05.

### IL-19 deficiency skews cytokine expression profiles toward Th17 cell expansion in macrophages

We then examined how IL-19 deficiency expands Th17 cells in the induction phase of EAE. First, we evaluated the mRNA expression level of IL-19 receptor (heterodimer of IL-20Rα and IL-20Rβ subunits) in the splenic immune cells such as macrophage, dendritic cell (DC), and CD4^+^ helper T cell. We found that both the IL-20Rα and IL-20Rβ subunits were more highly expressed in CD11b^+^ macrophages and CD4^+^ helper T cells than in CD11c^+^ DCs (Fig. S1). These results suggest that IL-19 mainly affects macrophages and CD4^+^ helper T cells.

Next, we assessed whether IL-19 directly differentiates naïve CD4^+^ T cells into Th17 cells. Naïve T cells were polarized using immobilized CD3 and CD28 antibodies in the presence of IL-6 and transforming growth factor β1 (TGF-β1), with or without IL-19. Quantitative PCR (qPCR) and flow cytometry revealed that IL-19 did not alter the differentiation of naïve T cells into Th17 cells (Fig. S2).

Because antigen-presenting cells (APCs) are crucial for differentiation of naïve T cells into effector T cells, we evaluated the expression levels of cytokines required for Th17 cell expansion in splenic macrophages and DCs. Interestingly, IL-19^−/−^ macrophages exhibited a significant increase in mRNA levels of the genes encoding IL-1β, IL-6, TGF-β, IL-12 p40, IL-23 p19, and tumor necrosis factor α (TNF-α), which play pivotal roles in Th17 cell differentiation and expansion (Fig. 4). Although a previous study showed that IL-19 increases IL-10 expression ^26^, our data showed that IL-19 deficiency did not alter the *Il10* mRNA level in macrophages (Fig. 4). By contrast, IL-19^−/−^ DCs did not exhibit a significant alteration in the expression levels of these cytokines (Fig. S3). These findings indicated that IL-19 deficiency skews the cytokine expression profiles toward Th17 cell differentiation and expansion in macrophages. Conversely, our data suggested that IL-19 suppresses Th17-skewed condition by activating macrophages.

**Figure 4.**
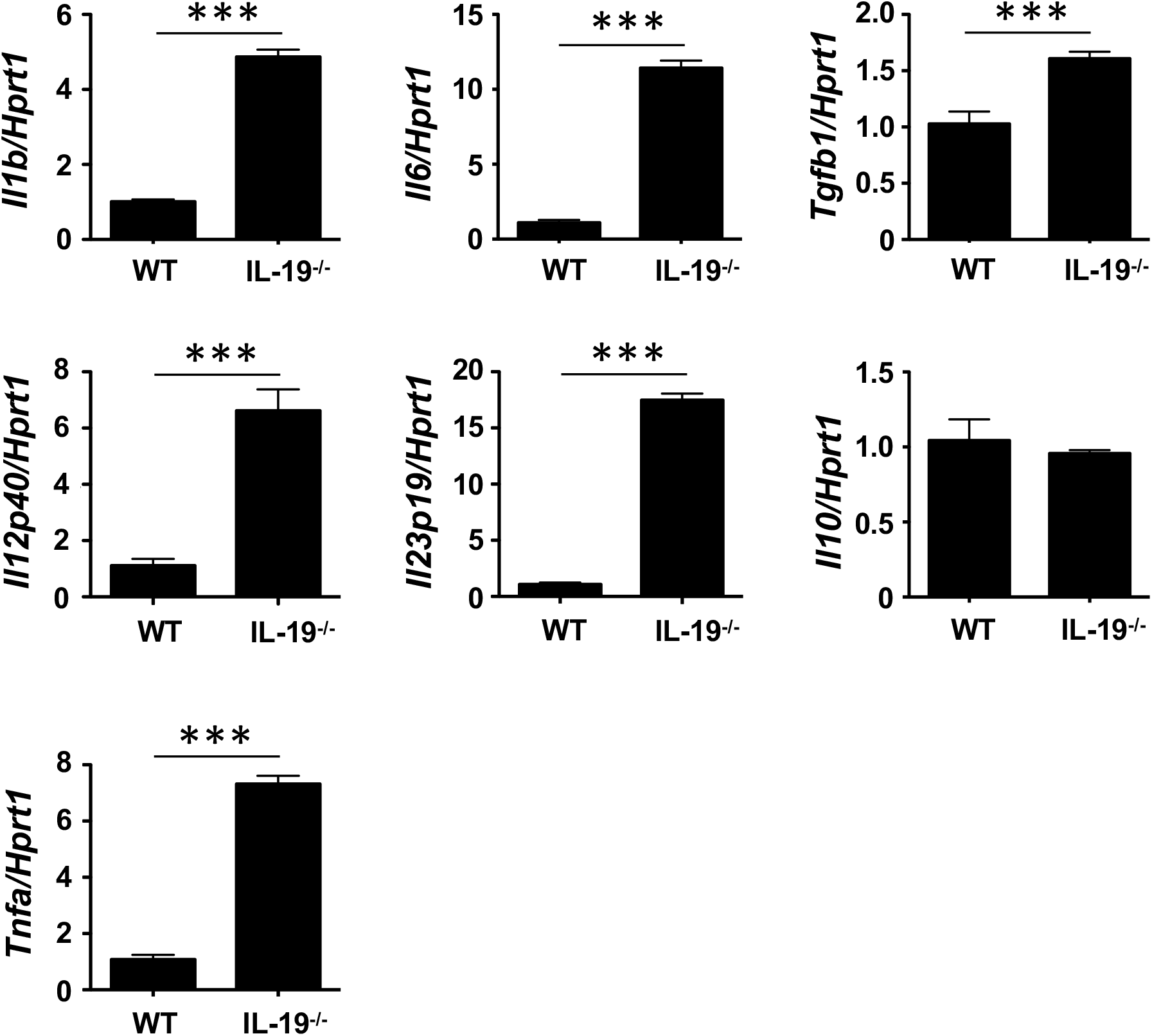
IL-19 deficiency upregulates Th17 cell differentiation–associated cytokines in macrophages. (A) qPCR data for mRNAs encoding IL-1β, IL-6, TGF-β1, IL-12 p40, IL-23 p19, IL-10, and TNF-α in splenic macrophages of EAE mice. Assessments were performed 7 days after immunization. Data represent means ± SD. ***, *p* < 0.0001 (n = 6).

### IL-19 deficiency promotes MHC class II expression in macrophages

To determine whether IL-19 signaling contributes to antigen presentation by macrophages, we assessed the expression of major histocompatibility complex (MHC) class II (H2-Ab) and co-stimulatory molecules (CD80 and CD86) in splenic CD11b^+^ macrophages from WT and IL-19^−/−^ EAE mice. (Fig. 5A). IL-19 deficiency significantly enhanced expression of the gene encoding MHC class II, whereas the genes encoding co-stimulatory molecules CD80 and CD86 were not affected (Fig. 5A). Flow cytometric data corroborated the enhanced presentation of MHC class II in IL-19^−/−^ splenic macrophages (Fig. 5B). These observations suggested that IL-19 also suppresses the antigen-presenting activity of macrophages.

**Figure 5.**
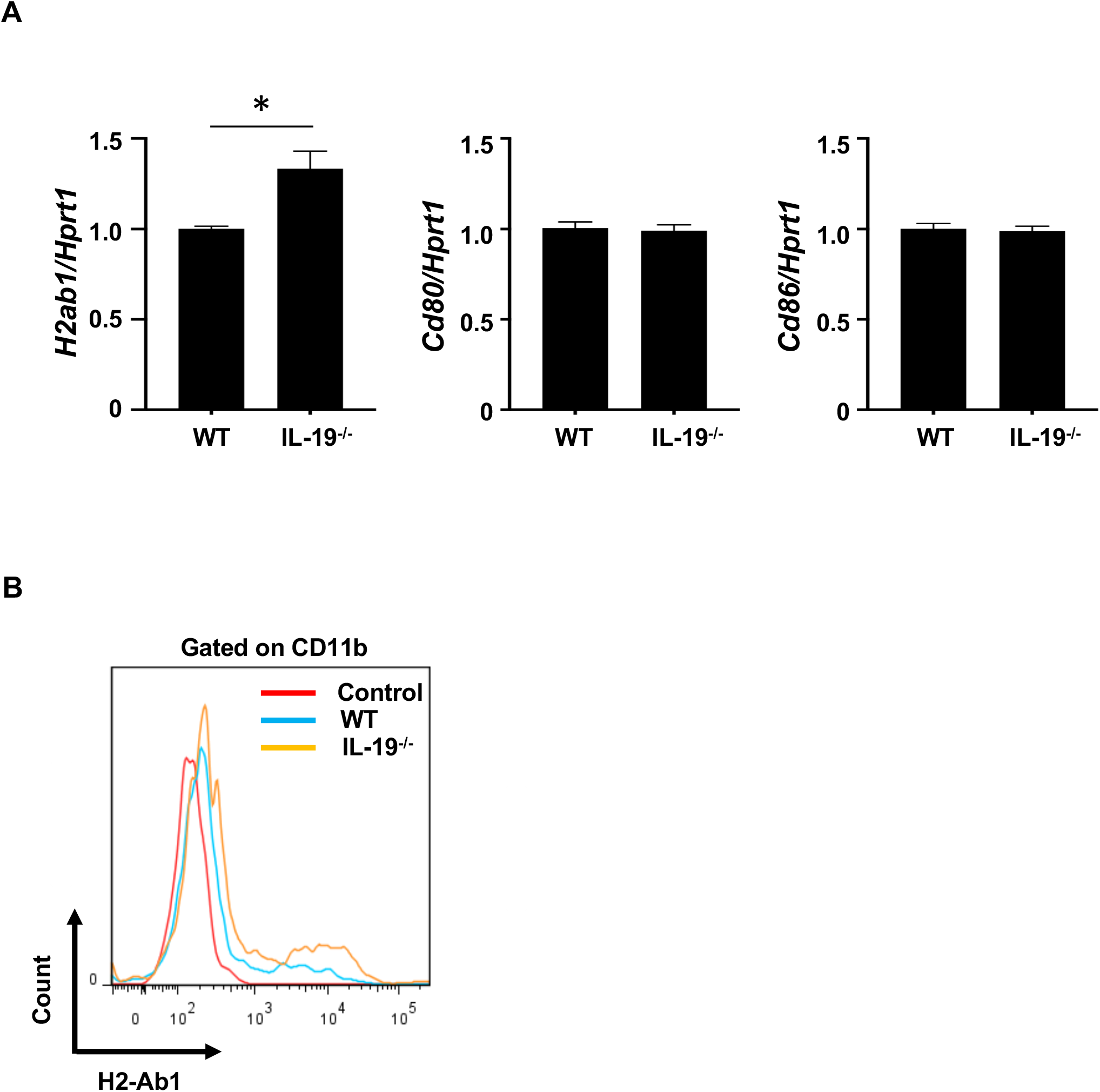
IL-19 deficiency enhances antigen-presenting activity in macrophages. (A) qPCR data for mRNAs encoding MHC class II (H2-Ab), CD80, and CD86 in the splenic macrophages on day 7 after immunization. Data represent means ± SD. *, *p* < 0.05 (n = 6). (B) Representative flow cytometric data for MHC class II (H2-Ab) presentation in splenic macrophages.

### Treatment with IL-19 abrogates EAE

To determine whether exogenous IL-19 abolishes the effect of IL-19 deficiency in EAE, we treated IL-19^−/−^ EAE mice with recombinant mouse IL-19 protein (20 ng/g of body weight) by intraperitoneal injection every other day, starting on day 2 post-immunization. As shown in Figure 6A, administration of IL-19 to IL-19^−/−^ mice abolished the exacerbation of EAE (Fig. 6A, IL-19^−/−^ + IL-19). We then investigated the therapeutic effect of IL-19 on EAE. When we treated WT EAE mice with recombinant mouse IL-19 protein in the same manner, we found that IL-19 treatment almost completely inhibited EAE (Fig. 6B). These results indicated that IL-19 represents a potential target for MS therapy.

**Figure 6.**
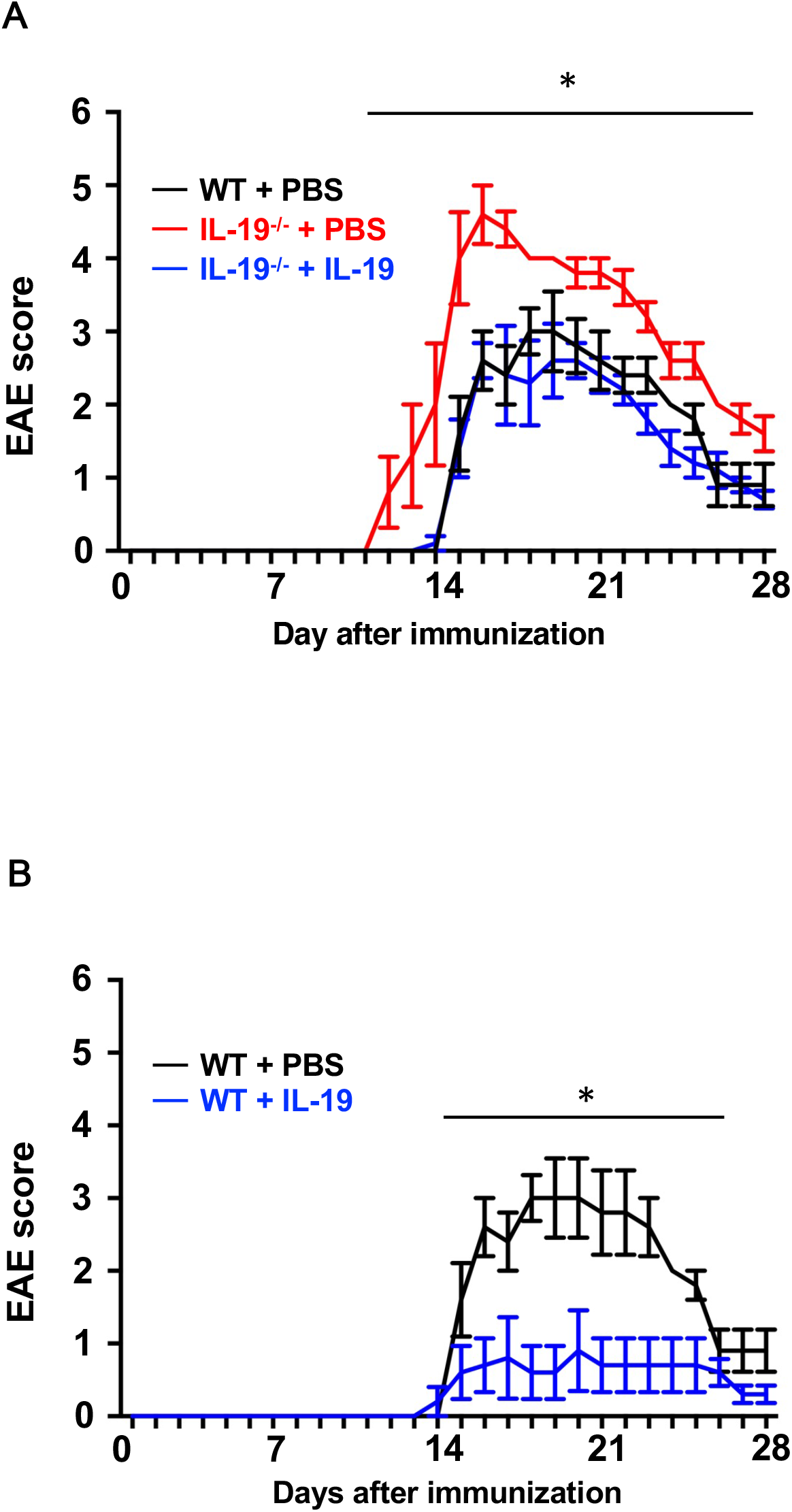
Treatment with recombinant IL-19 alleviates EAE. (A) EAE clinical score. WT + PBS (black): WT EAE mice treated with PBS; IL-19^−/−^ + PBS (red): IL-19^−/−^ EAE mice treated with PBS; IL-19^−/−^ + IL-19 (blue): IL-19^−/−^ EAE mice treated with IL-19. (B) EAE clinical score. WT + PBS (black), WT EAE mice treated with PBS; WT + IL-19 (blue), WT EAE mice treated with IL-19. Data represent means ± SD. *, *p* < 0.05 (n = 5).

## Discussion

In this study, we demonstrated that endogenous IL-19 negatively regulates development of EAE by inhibiting macrophage activation, and that IL-19 treatment effectively abrogates EAE. As shown in Fig. 1D and 1E, endogenous *Il19* mRNA expression was upregulated at EAE onset and downregulated at EAE peak in the spleen, whereas it was elevated at EAE peak in the CNS. We have previously identified IL-19 as a negative-feedback regulator to limit proinflammatory response of macrophages and microglia in autocrine/paracrine manners^11, 12^. From this point of view, these data imply that endogenous IL-19 increases to suppress inflammation accompanied by macrophage/microglia activation as disease progresses from the periphery to CNS, although it is insufficient to halt EAE progression.

Th17 cell infiltration in the CNS is considered critical for the development of EAE^5, 27^. Our data revealed that IL-19 deficiency increased CNS infiltration of Th17 cells, but not Th1 cells (Fig. 2). In addition, IL-19 deficiency enhanced the peripheral expansion of Th17 cells in the induction phase of EAE (Fig. 3). Conversely, our data indicated that IL-19 negatively regulates Th17 cell differentiation and expansion, which is critical for EAE development.

However, IL-19 did not directly affect differentiation of Th17 cells from naïve T cells (Fig. S2). APCs such as macrophages, microglia, and DC are also essential for effector T cell differentiation and expansion in EAE^28^. Specifically, IL-1β, IL-6, IL-23, TGF-β1, and TNF-α released by APCs play pivotal roles in Th17 cell differentiation and expansion^5, 29, 30, 31 32, 33^. In this study, we revealed that IL-19 deficiency significantly upregulated these Th17 cell differentiation–associated cytokines in macrophages, but not in DCs (Fig. 4 and Fig. S3). These phenomena were correlated with the expression level of IL-19 receptor (IL-20Rα and IL-20Rβ heterodimer), which was highly expressed in macrophages, but not in DCs or helper T cells (Fig. S1). Thus, IL-19 might negatively regulate Th17 cell differentiation and expansion through inhibition of cytokine release from macrophages.

Moreover, activated APCs highly express MHC class II and co-stimulatory molecules (CD80 and CD86) on the cell surface and acquire the ability to prime T cells^34^. As shown in Fig. 5, IL-19 deficiency further enhanced MHC class II expression in macrophages. Taken together, our findings suggest that IL-19 suppresses Th17 cell differentiation and expansion by suppressing cytokine production and antigen presentation in macrophages. Interestingly, a previous study reported that IL-17A induces IL-19 production^19^. Thus, IL-19 also might serve as a negative regulator of further Th17 cell polarization.

In addition, activated macrophages and microglia directly contribute to neuroinflammation-induced demyelination by releasing proinflammatory cytokines such as IL-1β; IL-6 and TNF-α^28, 35, 36, 37^. In our study, IL-19 deficiency increased the levels of these proinflammatory cytokines in macrophages and exacerbated EAE until the late phase of the disease. We previously revealed that IL-19 secreted from activated macrophages and microglia suppressed their proinflammatory responses in an autocrine/paracrine manner^11, 12^. Therefore, IL-19 might suppress development of EAE by dual inhibition of both autoreactive Th17 cell expansion and macrophage/microglia-mediated CNS neuroinflammation.

Although the IL-19 signaling pathway has not been fully elucidated, IL-19 mediates its downstream signaling at least by STAT3 activation^8, 9, 12, 38^. However, it remains controversial whether STAT3 activation is beneficial or harmful with regard to autoimmune-mediated neuroinflammation. Previous studies also reported that STAT3 activation in myeloid cells (including macrophages and microglia) exacerbates EAE^39, 40^. By contrast, STAT3 ablation worsens neuroinflammation in mice with spinal cord injury^41^, and STAT3 activation alleviates cuprizone-induced CNS demyelination^42^. This discordancy may depend on spatiotemporally specific activation of STAT3^43^. Indeed, in contrast to macrophages, IL-19 deficiency did not affect activation of DCs. Recent clinical trials of JAK/STAT inhibitors for autoimmune diseases have revealed a complicated signal network of cytokine/STAT axes in multiple cell types^44^. Further studies are needed to elucidate the precise IL-19 signaling pathway in each cell type.

In summary, we revealed that IL-19 deficiency exacerbated EAE by upregulating Th17 cell differentiation–associated cytokines and enhancing antigen presentation in macrophages, followed by Th17 cell expansion and infiltration in the CNS. We also demonstrated that IL-19 administration potently prevented development of EAE. Therefore, enhancement of IL-19 signaling represents a promising therapeutic strategy against MS and other Th17-mediated autoimmune diseases.

## Material and Methods

### Reagents

MOG peptide 35–55 (MOG_35–55_; MEVGWYRSPFSRVVHLYRNGK) was synthesized and purified by Operon Biotechnologies (Tokyo, Japan). Incomplete Freund’s adjuvant was obtained from Sigma-Aldrich (St. Louis, MO, USA). Heat-killed *Mycobacterium tuberculosis* H37Ra was obtained from Difco (Detroit, MI, USA), and pertussis toxin was obtained from List Biological Laboratories (Campbell, CA, USA). Recombinant mouse IL-6, IL-19, and TGF-β1 were obtained from R&D Systems (Minneapolis, MN, USA).

### Animals

All animal experiments were conducted under protocols approved by the Animal Experiment Committee of Nagoya University (approved numbers: 15017 and 15018) and Yokohama City University (approved number: F-A-19-036). C57BL/6J (B6) mice were purchased from Japan SLC (Hamamatsu, Japan). IL-19^−/−^ mice (B6 background) ^11, 12^ were obtained from Regeneron Pharmaceuticals (Tarrytown, NY, USA).

### EAE induction and treatment studies

MOG-EAE was induced as previously described ^45, 46, 47^. In brief, 8-week-old female mice were immunized subcutaneously at the base of the tail with 0.2 ml of emulsion containing 200 μg MOG_35–55_ in saline, combined with an equal volume of complete Freund’s adjuvant containing 300 mg heat-killed *Mycobacterium tuberculosis* H37Ra. The mice were intraperitoneally injected with 200 ng pertussis toxin on days 0 and 2 post-immunization. To investigate the effect of IL-19, EAE mice were treated with mouse recombinant IL-19 protein (20 ng/g of body weight) by intraperitoneal injection every other day starting on day 2 post-immunization, according to a modification of a previously reported method ^48, 49^. The mice were assessed daily for clinical signs of EAE, according to the following grading system: 0, normal; 1 - limp tail or mild hind limb weakness; 2 - moderate hind limb weakness or mild ataxia; 3 - moderate to severe hind limb weakness; 4 - severe hind limb weakness, mild forelimb weakness or moderate ataxia; 5 - paraplegia with moderate forelimb weakness; and 6 - paraplegia with severe forelimb weakness, severe ataxia, or moribundity.

### Isolation of cells from spleen and lumbar spinal cord

Mononuclear cells were collected from the spleen and lumbar spinal cord as described previously ^45, 46, 47^. CD4^+^, CD11b^+^, and CD11c^+^ cells were isolated using the MACS system (Miltenyi Biotec, Bergisch Gladbach, Germany). Helper T cell differentiation was induced as described previously ^5, 32, 47^. For flow cytometric analysis, cells were stained with PerCP/Cy5.5 or BV421-conjugated anti–mouse CD4 rat monoclonal antibody (RM4-5; BD Biosciences, Franklin Lakes, NJ, USA). The cells were then fixed and permeabilized with Cytofix/Cytoperm reagent (BD Biosciences) and stained with PE-conjugated anti–mouse IFN-γ rat monoclonal antibody (XMG1.2; BD Biosciences) and APC- or PE-conjugated anti–mouse IL-17A rat monoclonal antibody (TC11-18H10; BD Biosciences). The samples were analyzed using a FACS Aria III system (BD Biosciences) and the FlowJo software (FlowJo, Ashland, OR, USA).

### Histological analysis

Histological analysis was performed as previously described ^45^. Mice with peak EAE were anesthetized and perfused transcardially with 4% paraformaldehyde in 0.1 M PBS. Lumbosacral spinal cords were immediately removed, postfixed in 4% paraformaldehyde, and embedded in paraffin. Eight-micron-thick sections were stained with hematoxylin and eosin. Stained sections were analyzed on a NanoZoomer 2.0-RS slide scanner (Hamamatsu Photonics, Hamamatsu, Japan).

### RNA extraction and reverse-transcription polymerase chain reactions (RT-PCRs)

We evaluated the expression levels of proinflammatory factors in the lumbar spinal cords and spleens by qPCR as described previously ^12, 45^. In brief, lumbar spinal cords and spleens were collected from EAE mice at pre-immunization, EAE onset, and EAE peak (approximately on days 0, 10, 16 post-immunization, respectively). Total RNA was isolated with an RNeasy Mini Kit (Qiagen, Valencia, CA, USA) and reverse transcribed with SuperScript III (Life Technologies, Carlsbad, CA, USA). Expression levels of mRNAs were evaluated by qPCR using SYBR Select Master Mix (Applied Biosystems, Foster City, CA, USA) on a Rotor-Gene Q (Qiagen) or LightCycler 96 (Roche). Mouse gene–specific primers were obtained from Life Technologies (Table 1). Gene-expression values were determined using the ΔΔC_T_ method. Levels of mRNAs of interest were standardized to the geometric mean of the level of hypoxanthine phosphoribosyltransferase 1 (*Hprt1*). Assays were carried out in three independent trials.

**Table 1.**
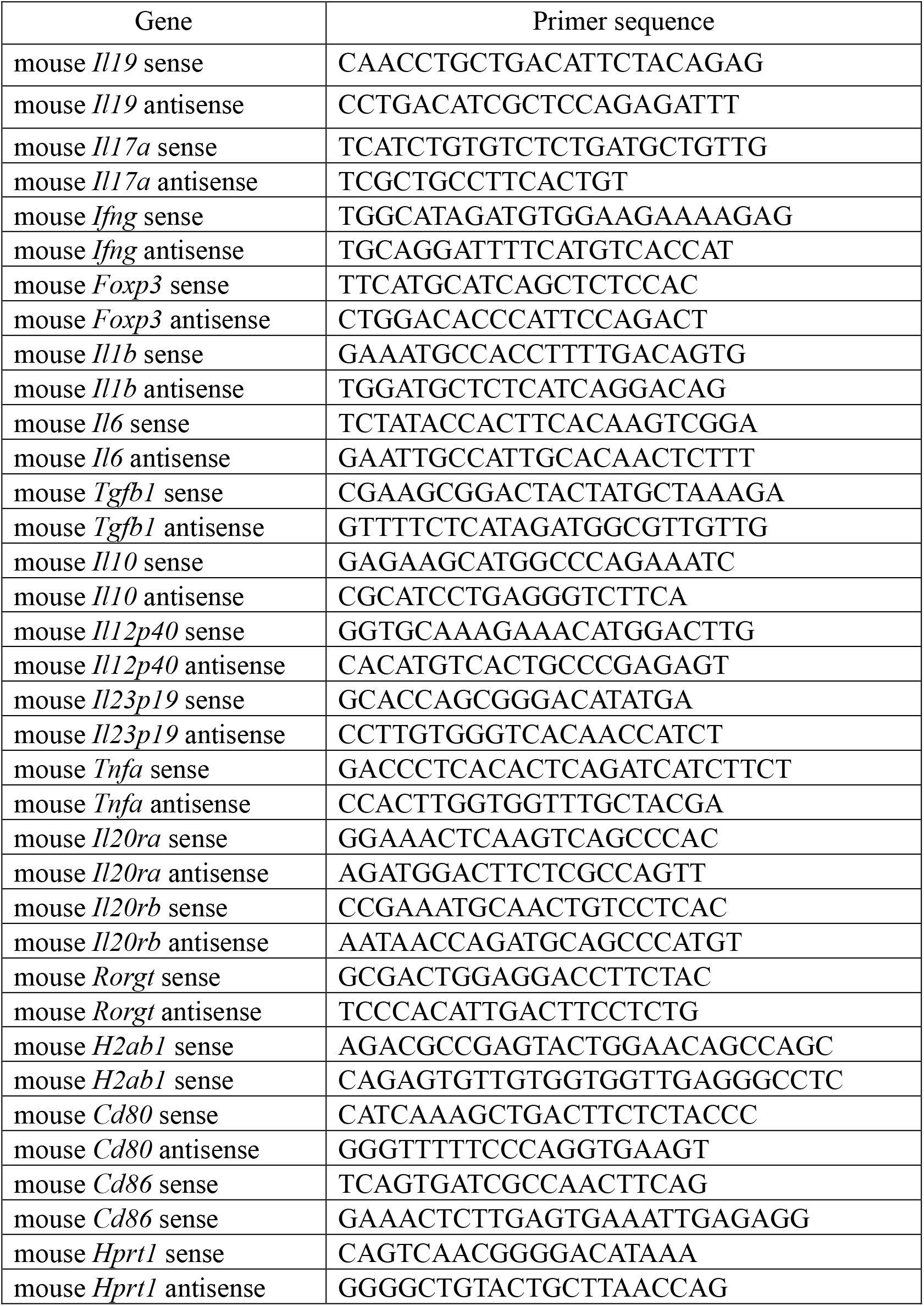
Primers for qPCR.

### Statistical analysis

Statistical significance was analyzed using Student’s t-test or one-way analysis of variance (ANOVA) followed by post hoc Tukey’s test in GraphPad Prism version 8 (GraphPad Software, La Jolla, CA, USA).

### Data availability

The datasets generated and analyzed during the study are available from the corresponding author upon reasonable request.

## Acknowledgements

This work was supported by grants-in-aid for Scientific Research from the Ministry of Education, Culture, Sports, Science and Technology of Japan (H.T.); grants from the Ministry of Health, Labour and Welfare of Japan (A.S., H.T.); the Program for Promotion of Fundamental Studies in Health Sciences of the National Institute of Biomedical Innovation (NIBIO) of Japan (A.S.); and a grant from the Naito Foundation (H.T.).

## Author contributions

H.H., B.P., A.S., and H.T. designed the research; H.H., B.P., H.K., Y.O., J.S., K.T., Y.A., and H.T. performed the research; H.H., B.P., H.K., F.T., A.S., and H.T. analyzed the data; and H.H., B.P., and H.T. wrote the paper.

## Supplementary Figure Legends

**Figure S1. IL-19 receptor heterodimer subunits IL-20Rα and IL-20Rβ are more highly expressed in macrophage and helper T cells than in dendritic cells.**

(A) qPCR for mRNA encoding IL-20Rα in CD11b^+^ macrophages, CD11c^+^ dendritic cells, and CD4^+^ helper T cells in the spleen. (B) qPCR data for mRNA encoding IL-20Rβ in CD11b^+^ macrophages, CD11c^+^ dendritic cells, and CD4^+^ T cells in the spleen. Data are represented as means ± SD. *, *p* < 0.05 (n = 3).

**Figure S2. IL-19 does not alter differentiation of naïve T cells into Th17 cells.**

(A) qPCR data for mRNAs encoding IL-17A and RORγt. (B) Representative flow cytometric data for IL-17A expression. (C) Quantitative analysis of (B). Data are represented as means ± SD. *, *p* < 0.05 (n = 5).

**Figure S3. IL-19 deficiency does not alter the expression levels of Th17 cell differentiation–associated cytokines in dendritic cells.**

qPCR data for mRNAs encoding IL-1β, IL-6, TGF-β1, IL-12 p40, IL-23 p19, IL-10, and TNF-α expression in splenic dendritic cells of EAE mice. Assessments were performed 7 days after immunization. Data are represented as means ± SD (n = 6).

