## Supplemental Figures for "Interleukin-19 alleviates experimental autoimmune encephalomyelitis by attenuating antigen-presenting cell activation"

### Supplementary Figure Legends

**Figure S1. IL-19 receptor heterodimer subunits IL-20Rα and IL-20Rβ are more highly expressed in macrophage and helper T cells than in dendritic cells.**

(A) qPCR for mRNA encoding IL-20Rα in CD11b$^+$ macrophages, CD11c$^+$ dendritic cells, and CD4$^+$ helper T cells in the spleen. (B) qPCR data for mRNA encoding IL-20Rβ in CD11b$^+$ macrophages, CD11c$^+$ dendritic cells, and CD4$^+$ T cells in the spleen. Data are represented as means ± SD. *, $p < 0.05$ (n = 3).

**Figure S2. IL-19 does not alter differentiation of naïve T cells into Th17 cells.**

(A) qPCR data for mRNAs encoding IL-17A and RORγt. (B) Representative flow cytometric data for IL-17A expression. (C) Quantitative analysis of (B). Data are represented as means ± SD. *, $p < 0.05$ (n = 5).

**Figure S3. IL-19 deficiency does not alter the expression levels of Th17 cell differentiation–associated cytokines in dendritic cells.**

qPCR data for mRNAs encoding IL-1β, IL-6, TGF-β1, IL-12 p40, IL-23 p19, IL-10, and TNF-α expression in splenic dendritic cells of EAE mice. Assessments were performed 7 days after immunization. Data are represented as means ± SD (n = 6).

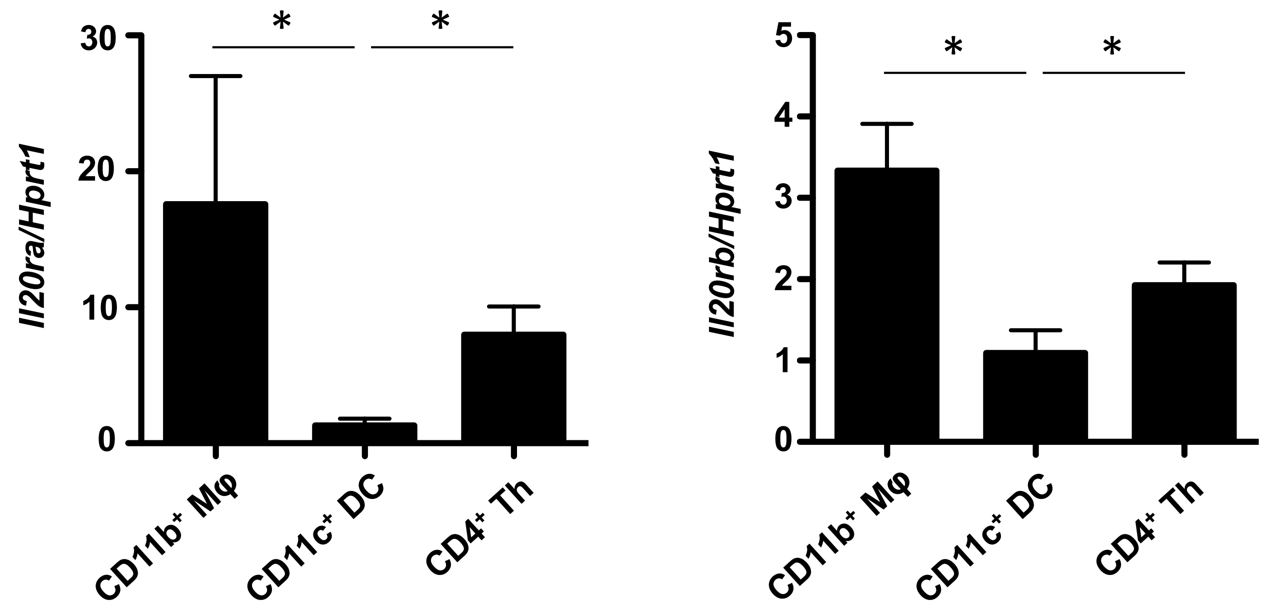


Horiuchi et al, Figure. S1

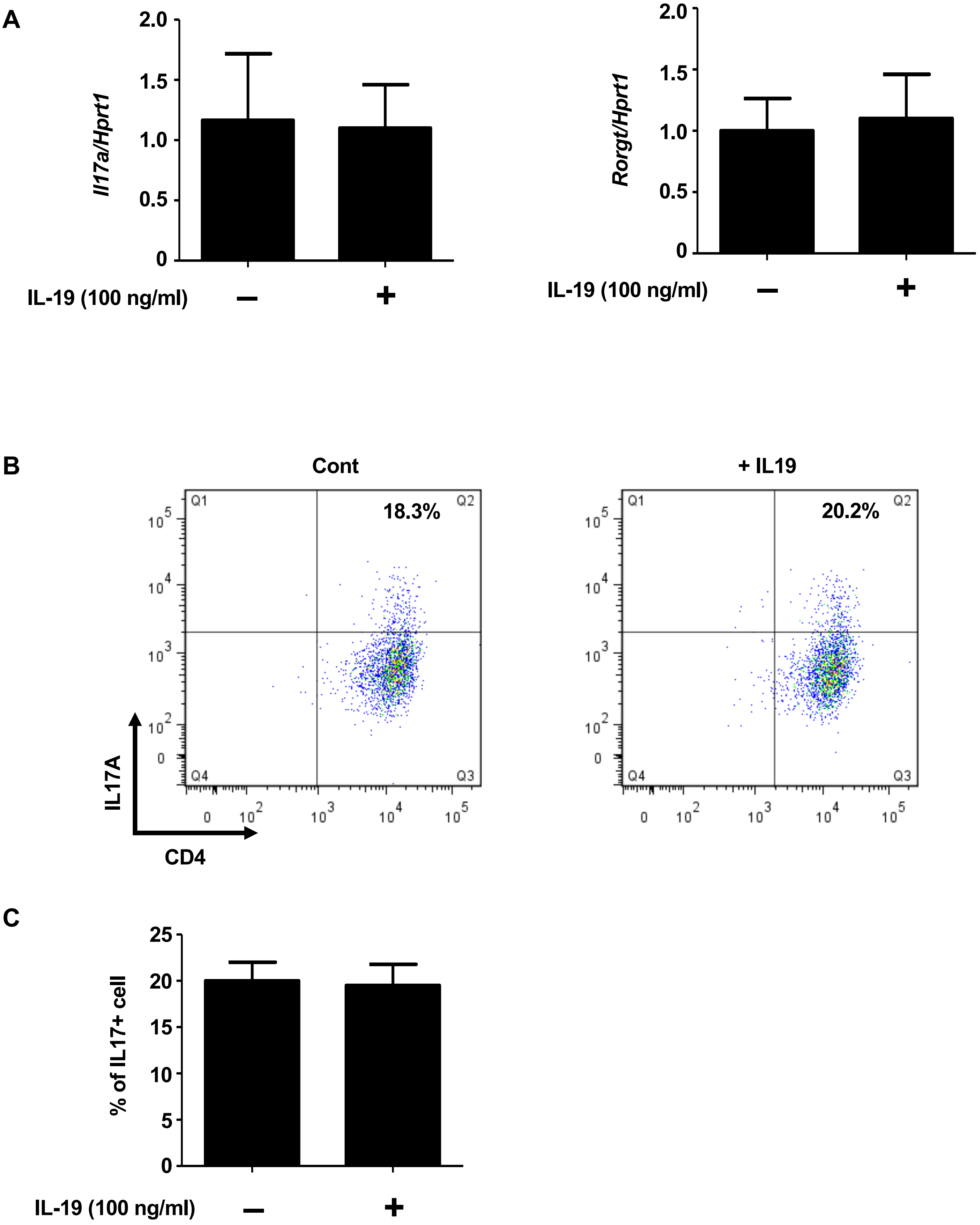


**Horiuchi et al, Figure. S2**

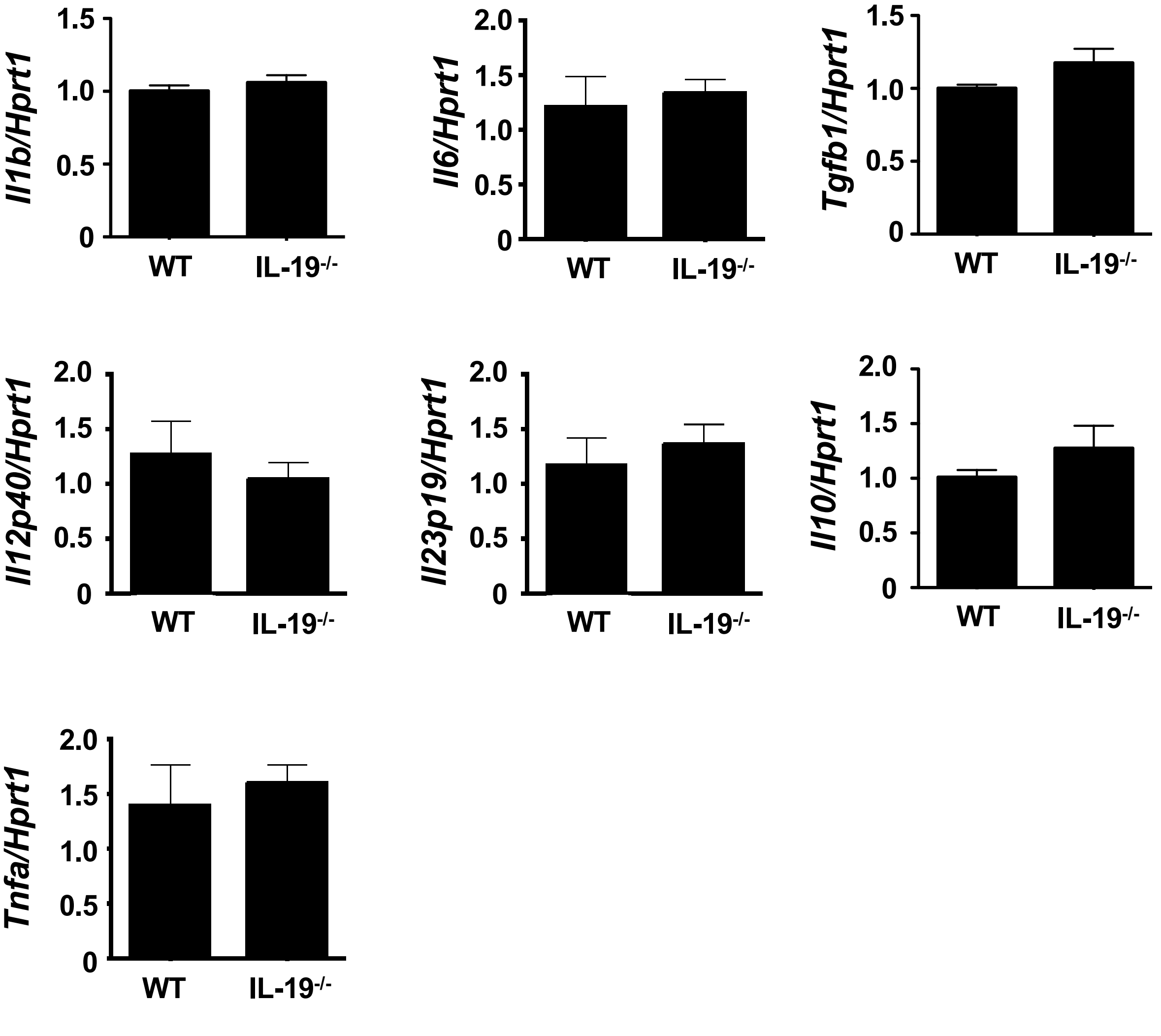


Horiuchi et al, Figure. S3
